# Open top multi sample dual view light sheet microscope for live imaging of large multicellular systems

**DOI:** 10.1101/2023.09.28.559925

**Authors:** F. Moos, S. Suppinger, G. de Medeiros, K. Oost, A. Boni, C. Rémy, S. Weevers, C. Tsiairis, P. Strnad, P. Liberali

**Author notes:** These authors contributed equally (F. Moos, S. Suppinger).

## Abstract

Multicellular systems grow over the course of weeks from single cells to tissues or even full organisms composed of several thousands of cells, making live imaging of such samples challenging. To bridge these wide spatiotemporal scales, we present an open top dual view and dual illumination light sheet microscope dedicated for live imaging of large specimens at single cell resolution. The novel configuration of objectives together with a flexible and customizable multi-well mounting system combines dual view with high throughput multi-position imaging. We use the newly developed microscope to image a wide variety of samples and highlight its capabilities to gain quantitative single-cell information in large specimens such as mature intestinal organoids and gastruloids.

## Main Text

Visualizing the dynamics of individual cells shaping complex tissues and understanding the underlying molecular mechanisms is an overarching goal in cell and developmental biology. However, these complex biological phenomena often cross large spatial and temporal biological scales, since multicellular systems can grow from a few micrometers in size to several hundred micrometers over the course of many days. Furthermore, biological processes and especially *in vitro* model systems are often affected by sample-to-sample heterogeneity. Designing a microscope for live imaging of such systems is challenging because it must provide high sample throughput within each experiment to draw conclusions despite this heterogeneity. Additionally, it must provide sufficient spatial- and temporal resolution and a high image quality for large light-scattering samples while minimizing light dosage and keeping the sample conveniently accessible to the user. Light sheet microscopy is one of the techniques which overcomes some of these challenges, due to its low phototoxicity and optical sectioning [1]–[3]. For large specimens, multi-view or SimView light-sheet microscopy has provided improved image quality by acquiring images from opposing directions using sample rotation or multiple objective lenses [4]–[7]. These techniques are however limited to imaging of either just one or very few [8] samples per experiment and do not provide convenient multi-well sample mounting. On the other hand, open top [9], oblique plane [10], SCAPE [11] or inverted [12]–[14] light sheet microscopes have been developed to enable multi sample imaging, in which the sample is mechanically supported from the bottom by a custom sample holder [12] or by standard culture dishes [15] while granting direct accessibility from the top allowing easy sample mounting. However, these systems lack the possibility for multi-view imaging from opposing detection sides and are therefore not well suited for larger specimens.

Here we present an open top, dual view, and dual illumination light sheet microscope, combining the advantages of multi-view light sheet microscopy with an open top geometry and a multi-well sample holder enabling long term multi position 3D live imaging of large multicellular systems. We show the capabilities of the system to achieve high image quality in a variety of model systems by long term live imaging of murine intestinal, liver, and salivary gland organoids, gastruloids, *Hydra*, and human colon cancer organoids, reaching sizes of up to 550 µm and recordings for up to 12 days. Furthermore, we obtain quantitative features across biological scales and present a detailed single cell analysis through tracking and segmentation for intestinal organoids and gastruloids, which is only possible with the newly developed microscope.

Our light sheet microscope contains two opposing illumination objectives (Nikon 10X, NA 0.2) each tilted slightly upwards from the horizontal plane, illuminating the sample from two sides, and two opposing detection objectives imaging from two directions (Nikon 16X, NA 0.8, the system is also compatible with Nikon 25X, NA 1.1) (Fig. 1a-e and Extended Data 1). This geometry creates an unobstructed linear space just above the two illumination objectives (Fig. 1b, e) for a custom designed multi-well sample holder containing an array of up to four interchangeable sample chambers (wells) for multi-position imaging (Fig. 1f). Immersion medium (water) is placed in a reservoir filling the space between the two water immersion detection objectives. To obtain two opposing light sheets illuminating the sample at a largest possible angle we used super-long working distance air objectives, coupled the illumination light into the immersion medium through a glass window and designed a custom correction triplet lens compensating the spherical and chromatic aberrations. The objective area is temperature controlled and the sample is enclosed in a compartment with controlled CO2 concentration. An additional beam path is using one of the detection objectives as a condenser to illuminate the sample and acquire transmitted light images. To optimally mount different biological samples, we developed customizable chambers produced from fluoroethylene propylene (FEP) foils in a thermoforming process [16] (Fig. 1f and Methods). The chambers fit into the 6 mm space in between the two detection objectives and allow access for pipetting from the top and meeting the requirements of different biological samples. For specimens embedded in a matrix such as Matrigel, we developed chambers with a straight bottom (Fig. 1f) with small width to minimize degradation of image quality caused by Matrigel. For samples grown in suspension such as *Hydra* or gastruloids, we designed chambers with pocket sizes adapted to the size of the specimens. Since the sample holder and the molds for the imaging chambers can be produced easily with 3D printing or aluminum milling, a wide range of different geometries is possible, giving maximal flexibility. In this way, we can ensure growth and environmental conditions similar to experiments performed in standard plate formats providing consistency between microscopy data and other complementary experiments.

**Figure 1.**
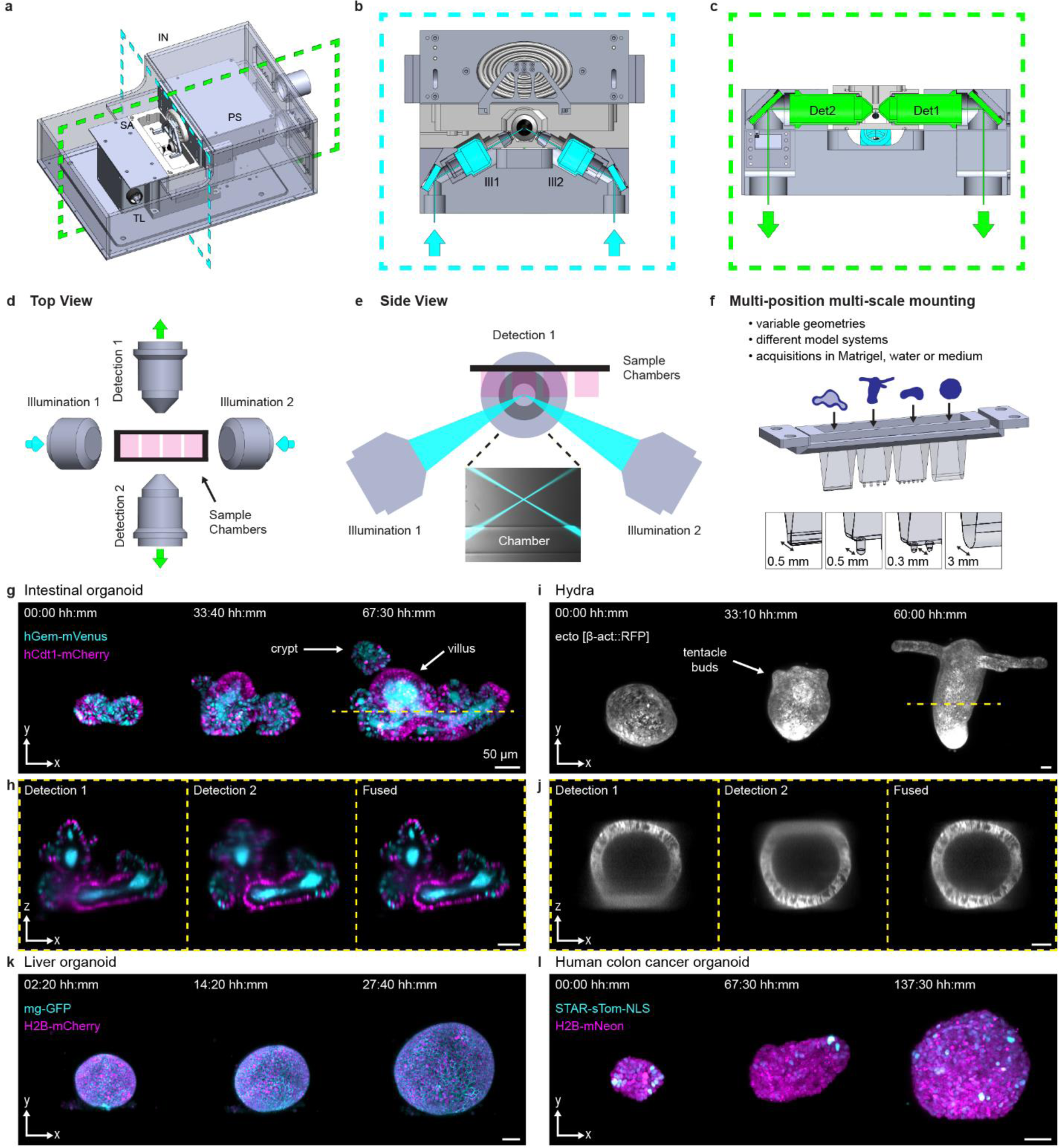
Open top dual view light sheet microscope with examples of time lapse acquisitions of various multicellular systems. a) Model of the dual view light sheet microscope, showing the incubator (IN), sample mounting area (SA), transmitted light (TL) and sample positioning unit (PS). Blue and green dashed lines indicate the positions of the cross sections shown in b) and c). b) Cross-section of the model of the microscope showing the arrangement of illumination objectives (Ill1, Ill2) highlighted in blue. c) Cross-section of the model of the microscope showing the arrangement of the two detection objectives (Det1, Det2) highlighted in green. d) Sketch of the top view of the microscope showing the arrangement of the two detection (Detection 1 and 2) and two illumination objectives (Illumination 1 and 2) with space for sample chambers. The green arrows indicate the direction of the emission light, and the blue arrows indicate the excitation light. e) Sketch of the side view of the objective arrangement. The illumination objectives are tilted to make space for the sample chambers. The zoom-in shows an image from the microscope with the illumination beams and the bottom of a sample chamber. f) Model of the sample holder with four sample chambers. Sample chamber geometries are adapted to the sample requirements such as their size or surrounding medium (Matrigel, medium and water). g) Maximum intensity projections (MIPs) along Z axis showing three time points from time-lapse acquisition with organoids expressing the Fucci2-reporter (hGem-mVenus and hCdt1-mCherry). The yellow line indicates the position of the cross section in h). Scale bar 50 µm. h) Cross section in XZ plane of the intestinal organoid shown in g) using Detection 1, Detection 2 and the fused data from both objectives. Scale bar 50 µm. i) MIPs along Z axis showing three time points from time-lapse acquisition of *Hydra* expressing an ectoderm-reporter (ecto [β-act::RFP]). The yellow line indicates the position of the cross section in j). Scale bar 50 µm. j) Cross section in XZ plane of the *Hydra* used shown in i) Detection 1, Detection 2 or the fused data from both objectives. Scale bar 50 µm. k) MIPs along Z axis showing three time points from a time-lapse acquisition of liver organoids expressing mg-GFP and H2B-mCherry reporter. Scale bar 50 µm. l) MIPs along Z axis showing three time points from time-lapse acquisition of human colon cancer organoids expressing STAR-sTom-NLS and H2B-mNeon. Scale bar 50 µm.

In our former work [17], we employed the predecessor of the here presented microscope [13] built with only one detection objective to track cells in developing intestinal organoids. However, the imaging depth provided was not sufficient to track cells in larger specimens, including mature organoids. The newly devised dual detection, dual illumination approach overcomes this hurdle. We imaged crypt and villus formation of maturing mouse intestinal organoids over the course of 3 days (Fig. 1g and Supplementary Video 1), using the cell cycle reporter, FUCCI2 [18]. The cell-cycle status was monitored in the context of crypt and villus formation. Using multi-position imaging, we acquired datasets for multiple organoids simultaneously. Dual-color imaging with single cell resolution within a depth of 360 µm with a temporal resolution of 10 minutes (Supplementary Video 1), allowed the visualization of the *in toto* dynamics of developing mouse intestinal organoids in unmatched detail. Visualizing single cells throughout the whole sample volume requires high image quality achieved through dual detection. We illustrate this by comparing the XZ sections of the individual detection objectives to the fused data (Fig. 1h and Methods). The sections of the single views show an expected degradation with increasing imaging depth, whereas the fused data is composed of optimal quality combined from both views and enabled the mapping of individual cells throughout the entire volume (Supplementary Video 2). This strategy is also necessary to image entire animals such as the cnidarian *Hydra* in 3D. We recorded *Hydra* regeneration for 2.5 days starting from newly formed spheroids cut from adult animals, which was previously shown to be driven by mechanical oscillations [19]. Imaging several individuals allowed the parallel observation of body axis formation along with the development of the foot and head structures such as the tentacles. (Fig. 1i, j and Supplementary Video 3 and 4). To the best of our knowledge, we are the first to show time lapse imaging of *Hydra* with this time resolution. Previously published work achieved a time resolution of 40 mins [20].

To further assess the improvement in image quality by using two opposing detection objectives, we compared the image quality with increasing imaging depth by calculating the Shannon Entropy of the Discrete Cosine Transform (DCT) [21] for each z-section of both detection objectives individually and the fused data. As example we performed this comparison with an intestinal and human colon cancer organoid (Extended Data Fig 2a-d). Both model systems show a clear decrease in image quality with increasing imaging depth and distance from the detection objective (Extend ended Data Fig 2b,d), which can be compensated by combining the data from two opposing objectives. Additionally, we compared the image quality of our developed system with another open-top light sheet microscope published in [13]. This system offers dual illumination (Nikon 10X, NA 0.3) and one detection objective (Nikon 25X, NA 1.1). We imaged the same gastruloid stained with the nuclear marker DRAQ5 in both systems (Extended Data Fig 2e,g) and evaluated the image quality by calculating Shannon Entropy of the DCT. The single-detection system shows a degrading image quality score with increasing imaging depth (Extended Data Fig 2f), whereas our microscope enables a higher image quality throughout the sample due to the opposing detection objectives (Extended Data Fig 2h). Together, these results show the importance of dual detection to guarantee high image quality in large specimens.

**Figure 2.**
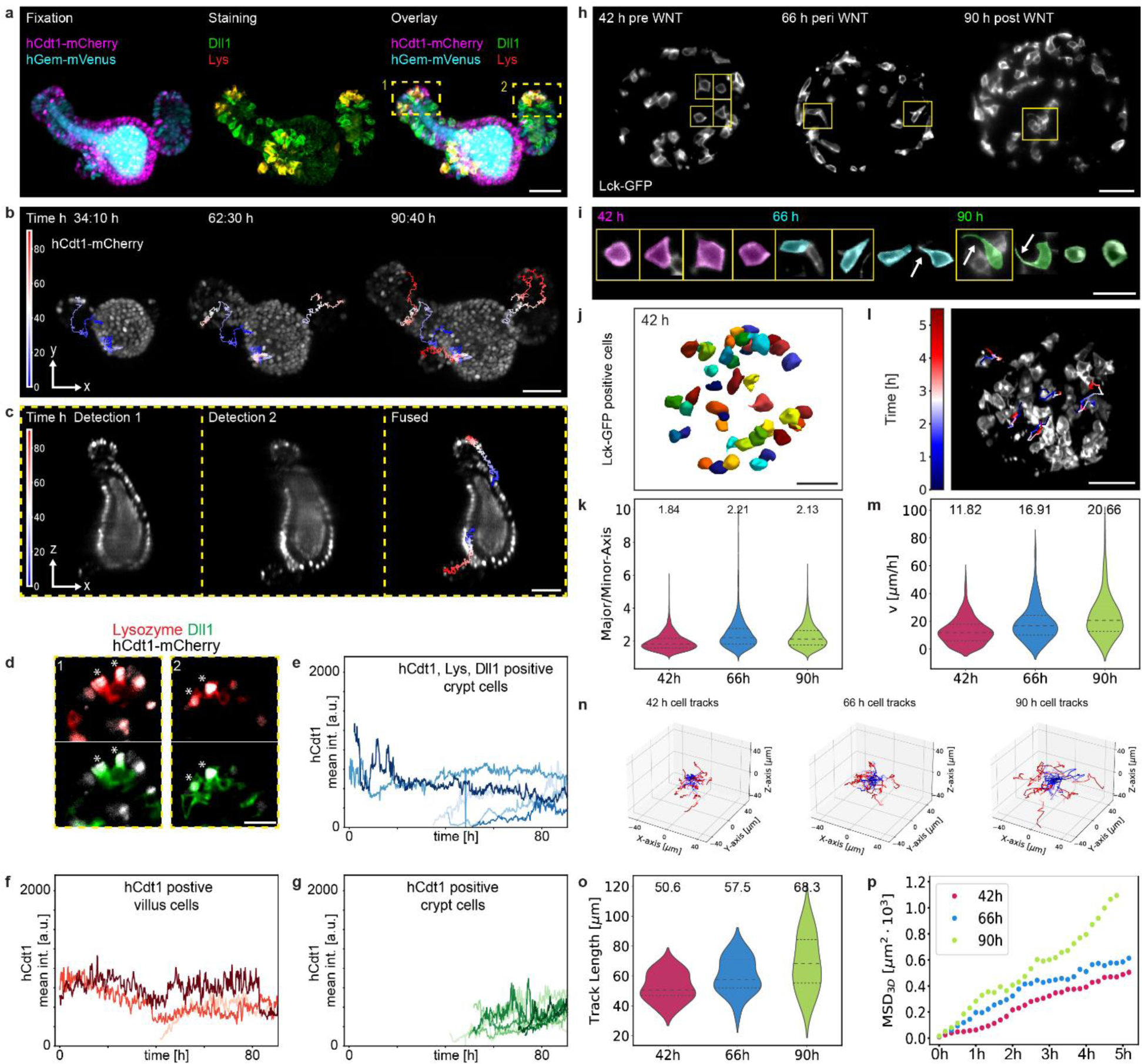
Single cell analysis of intestinal organoids and gastruloids. a) Intestinal organoid expressing hGem-mVenus and hCdt1-mCherry (left) was fixed and stained with antibodies specific for Lysozyme (Lys) and DLL1 (middle) at the last time point of a time-lapse acquisition (90:40 h). Overlay comparing the spatial localization of the staining with the original signal (right). Scale bar 50 µm. b) Maximum intensity projections of an intestinal organoid expressing hCdt1-mCherry at three different time points of a time lapse acquisition. Overlaid are the tracks of back-tracked cells color-coded over time. Scale bar 50 µm. c) Z-Projections of the intestinal organoid shown in b) of Detection 1, Detection 2 and the fused data. Overlaid are two representative tracks from back-tracked cells. Tracks are color coded for temporal progression. Scale bar 50 µm. d) Zoom-ins of the highlighted crypts of the intestinal organoid shown in a) with stars indicating triple positive cells for hCdt1, Dll1 and Lysozyme. Scale bar 50 µm. e) Quantification of hCdt1 intensity over time for cells that are triple positive (hCdt1, Dll1 and Lysozyme) and are located in the crypt at the last time point. Each color represents a single cell ((n = 5). f) Quantification of hCdt1 intensity over time for hCdt1-positive cells, that are located in the villus at the last time point. Each color represents a single cell (n = 3). g) Quantification of hCdt1 intensity over time for hCdt1-positive cells, that are located in the crypt at the last time point. Each color represents a single cell (n = 8). h) Z-planes of gastruloids expressing Lck-GFP at three different developmental stages (42h pre Wnt, 66h peri Wnt and 90h post Wnt). Yellow boxes highlight individual cells are shown in i). Scale bar 50 µm. i) Individual cells partly highlighted in h) from gastruloids 42h, 66h and 90 h after cell seeding. Arrows point towards cell protrusions. Scale bar 20µm. j) Example of 3D segmentation (Cellpose) of Lck-GFP positive cells of gastruloid 42h after cells seeding. Scale bar 50 µm. k) Comparison of the ratio of major and minor axis 42h, 66h aI nd 90 h after cell seeding, showing the median (values depicted in figure) and the first and third quartile. Measurements were performed over 33 timepoints on the 3D volumes of 3 gastruloids per imaging window (42 h, 66 h or 90 h). Summing up individual timepoints, a total of 4466 datapoints (42 h), 5723 datapoints (66 h) and 4598 datapoints (90 h) were analyzed. l) Maximum intensity projection of a gastruloid overlaid with the tracked cells over time. Scale bar 50 µm. m) Violin plots of cell velocity (µm/h) grouped by observation windows, showing the median (values depicted in figure) and the first and third quartile. The following numbers of datapoints (from 3 individual gastruloids) were tracked per observation window: n = 768 (42 h), n = 608 (66 h), n = 576 (90 h). n) Cell tracks of individual cells of gastruloids imaged for 5.5 h mounted at 42 h, 66 h and 90 h post seeding centered at the origin of the coordinate system, color coded for the temporal progression. Tracks were generated for 24 cells over 32 timepoints (42 h), 19 cells over 33 timepoints (66 h) and 18 cells over 32 timepoints (90 h). o) Distance plot displaying track length per cell for the individual imaging windows of Lck-GFP chimera gastruloids. Violin plots of cell velocity (µm/h) grouped by observation windows, showing the median (values shown in figure) and the first and third quartile. The track length was measured for 24 cells (42 h), 19 cells (66 h) and 18 cells (90 h). p) Mean square displacement averaged over all cells for the individual imaging windows of Lck-GFP chimera gastruloids. MSD measurements were performed over 32 timepoints for 24 cells (42 h), 19 cells (66 h) and 18 cells (90 h).

To illustrate the versatility of our system, we imaged a variety of samples from 200 µm to 550 µm in size and for up to 12 days of continuous imaging (see Supplementary Table 1 for imaging details of all acquisitions): murine liver organoids (Fig. 1k and Supplementary Video 5), human colon cancer organoids [22] (Fig. 1l and Supplementary Video 6 and 10), murine parotid salivary gland organoids (Extended Data Fig. 2i and Supplementary Video 7) and gastruloids (Extended Data Fig. 2j and Supplementary Video 8). In addition, the newly devised sample mounting strategy not only supports the development of various different specimens but also enables experiments with parallel chemical perturbations. During the same imaging experiment, each of the four chambers can be used for a particular condition with many individual positions imaged within each chamber. We conducted an analysis of the mechano-osmotic effects of prostaglandin E2 (PGE) [23], [24], Forskolin (a CFTR activator) [25], and a hyperosmotic shock induced by NaCl on intestinal organoids. The high temporal resolution of 3 minutes (Supplementary Video 9) revealed rapid organoid inflation and contraction in response to the respective treatments. Furthermore, to demonstrate the multi-positioning capabilities of our microscope we imaged in parallel 25 mature intestinal organoids with a time resolution of 10 mins covering a volume of 360 µm for each organoid (Supplementary Video 10).

After establishing that the microscope enables both long term (e.g. developing cancer organoids for up to 12 days, see Supplementary Video 11), highly dynamic imaging (e.g. perturbations of intestinal organoids with high temporal resolution (Supplementary Video 9)) while allowing multi-position imaging (Supplementary Video 10), we characterized its capabilities to obtain high quality single cell resolution data. Therefore, we performed live imaging of intestinal organoids expressing the cell cycle reporter FUCCI2 [18]. The above presented chamber design also allows the fixation of samples after live imaging. Leveraging this feature, we gained further insights into the cell type composition of formed organoids by combining dynamic information of fluorescent reporters with end-point functional readouts via antibody staining (Fig 2a), as proposed in [17]. Due to minimal movement of the sample even after the removal from the microscope for immunofluorescent staining, we imaged the re-mounted sample and overlaid the last live imaging timepoint with the stained organoid via 3D registration (Extended Data Fig. 2k and Methods). Here the Paneth cell marker Lysozyme (Lys) and the secretory cell marker Dll1 were used. This approach facilitated the detection, annotation and tracking of individual cells of interest. Newly identified triple positive cells (hCdt1, Lys and Dll1) were tracked back in time in order to monitor Paneth cell maturation and their cell cycle arrest in G0/G1. As expected, the initial position of the maturing Paneth cells was predictive of the eventual position of the organoid crypt (Fig 2b). This observation was made possible through dual illumination and dual detection imaging as cells moved to imaging depths where single cell resolution could not have been achieved with single detection only (Fig. 2c). The additional information gained from immunofluorescence allowed us to compare hCdt1^+^/Lys^+^/Dll^+^ Paneth cells, with hCdt1^+^/Lys^-^/Dll^-^ cells in the crypt (predominantly intestinal stem cells) and the villus of the organoid (mostly enterocytes) (Fig 2d). We further discovered that some of the enterocytes and Paneth cells were already terminally differentiated before the recording began, indicating longer lifespans. In contrast, other cells, especially in the crypt, were identified at the moment of division, allowing us to track their maturation with an average cell cycle length of 34.5 h at the time of fixation. Assessing the timing of emergence of each population it became apparent that Paneth cells emerge earlier than hCdt1 single positive cells in the crypt (most likely stem cells) and cells in the villus (enterocytes), suggesting a cell type specific cell cycle length and order of emergence (Fig. 2e-g). These findings highlight the significance of the newly devised imaging strategy, enabling us to discern cell type-specific dynamics and temporal relationships within the organoid. Such insights into cellular behavior and maturation processes would not have been attainable without the combination of live imaging and immunofluorescence within the entire organoid volume.

Unlike most other samples we imaged, gastruloids are dense structures aggregated from mouse embryonic stem cells [26] and thus are challenging to image with single cell resolution. Hence, we made use of this model system, to display the resolving capability of our light sheet microscope to achieve single cell resolution and to analyze the motility and shape features of individual cells in developing gastruloids. Standard gastruloid protocols utilize a Wnt activation pulse (Chiron99021, Chir) [26], [27] to increase mesoderm formation efficiency. This Wnt agonist treatment and the consequent activation of mesodermal programs potentially induce an epithelial to mesenchymal transition (EMT) like behavior and an increase in cell migration [28]. To analyze the cell shape of individual cells within dense cell populations, we generated chimeric gastruloids (Fig. 2h) expressing a membrane reporter (Lck-GFP) only in a subset of the cells (∼10%). Subsequently, we recorded the live dynamics of gastruloids before (42 h after aggregation), during (66 h after aggregation) and after the Wnt pulse (90 h after aggregation) for 5.5 hours with a 10 min time resolution (Supplementary Video 12). Additionally, we embedded the gastruloids in 40% Matrigel to prevent mechanical rotations in the sample chambers. 3D segmentation of single cells using Cellpose [29] (see Fig 2j) allowed the calculation of major/minor axis ratios, showing an increase in elongation of cells over time peaking during Chir treatment (Fig. 2k). Notably, we observed significant changes in cellular shape over the course of gastruloid development. During Wnt activation (time window imaged 66 h after aggregation) cells started to adopt a more elongated geometry that was then partially reverted in the time window imaged 90 h after aggregation. Additionally, at later stages (90 h after aggregation) a subpopulation of cells displayed long cell protrusions (Fig. 2i). This observation led to the hypothesis that cell motility is increasing along gastruloid development during the Wnt pulse. Using the Fiji plugin Mastodon [30] we tracked single Lck-GFP positive cells (Fig. 2l and 2m) and found that the median velocity of migration (µm/h) increased across the three chosen time windows. The strongest acceleration occurred during the Wnt pulse exhibiting a 1.43 fold increase in median speed compared to the time window before the Wnt pulse (Fig 2n). Moreover, the visualization of the individual cell tracks in 3D revealed that cells migrated over longer distances. Gastruloids imaged post Wnt activation (90 h after aggregation) also stood out, compared to the other assessed time windows, when looking at the travelled distance (track length) over the 5.5 h of imaging (Fig 2o). Evaluation of the 3D mean square displacement (MSD3D) suggested an increased speed and a change in migration behavior of cells tracked from 90 h onwards (Fig 2p). Together, these findings demonstrate an increase in cell migration over time, suggesting that Wnt activation not only induces the emergence of germ layer derivatives but also enhances cell motility and potentially coordinated migratory behavior in gastruloids [31]. While cells display pronounced elongation during the process, cell speed peaks after Wnt activation. The observed changes in cell shape, coupled with increased motility, suggest at least a partial EMT-like transition in gastruloids [26]. This trend remains consistent between gastruloids embedded in Matrigel and gastruloids grown in suspension (Extended Data Fig. 2l,m). Overall, these results demonstrate that this microscope provides data suitable for single-cell quantifications, even in challenging dense samples like gastruloids.

In summary, we present a dual view and dual illumination open top light-sheet microscope suitable for long term multi-position imaging of a wide range of large biological model systems (e.g. intestinal organoids, *Hydra*, gastruloids) at single cell resolution and with quality suitable for cell segmentation and tracking of cells in the entire organoid (Extended Data Fig 3a). The unique objective configuration enables the use of multi-well sample holders to monitor multiple experimental conditions simultaneously. Sample specific and flexible mounting was achieved by employing a thermoforming process to manufacture sample holders of various shapes. Unlike other microscopes [32], the production of such custom sample holders is simple and only requires minimal equipment.

Previously, different variations of open top light sheet microscopes have been developed to combine the 3D imaging capabilities of light sheet microscopy with multi-well plates or glass slides using either an single objective approach [10], [11], [33]–[36] (Extended Data Fig 3b,c) or objectives placed under the sample holder facing upwards [9], [12], [37] (Extended Data Fig 4d). Another solution proposed the use of two opposing illumination objectives to minimize shadowing effects [13] (Extended Data Fig 3f) and one detection objective. This solution still allowed the use of custom designed multi-well holders and multi-position imaging with access from the top. All these proposed solutions miss the possibility to image the sample from two opposing sides, which is crucial to visualize single cell dynamics within the whole specimen. On the contrary, multi-view light sheet microscopy was developed to provide high image quality throughout the sample by imaging the samples from multiple sides using several detection objectives (Extended Data Fig 3g) [4]–[6], [38]. However, these approaches are limited in sample throughput due to the constraint geometry of the sample holder or require the sample to be embedded, which is not suitable for every model system. Our system combines the advantages of the open top geometry with the multi-view approach (Extended Data Fig 3 h). Unlike other open top light sheet systems, the new dual-detection design enables high image quality throughout the entire sample allowing us to track and segment single cells in gastruloids.

In the future, a light sheet microscope with this objective configuration will be a promising platform for further technical advancements, such as precise perturbations at the single cell level using laser ablation or optogenetic stimulations. Integration of adaptive optics or imaging of optically cleared specimens could further enhance image quality. In conclusion, the microscope presented in this study enables dual view dual illumination multi position long-term imaging of large samples mounted in easily customizable open top multi-well sample holder, features which are all required to conduct live imaging experiments of multicellular systems.

## Acknowledgment

We thank Markus Klement (Department of Biosystems Science and Engineering (DBSSE)) for producing sample holders and sample chamber molds, Enrico Tagliavini for IT support. We acknowledge Véronique Kalck for general support in the lab, organoid cultivation and genotyping of the newly established mouse strains. Chiara Azzi and Marietta Hartl contributed to the efforts of mESC line engineering by providing plasmids and performing parts of the cloning work. This project received funding from the SNSF Sinergia grant (CRSII5_189956) and the European Research Council under the European Union’s Horizon 2020 research and innovation program (grant agreement number 758617).

## Author contributions

P.L. and P.S. supervised the work, P.L., P.S. and A.B. originally conceived the project. F.M., G.d.M., A.B., C.R. and P.S. designed and constructed the microscope. P.S. wrote the microscope software. F.M. performed experiments and developed the data analysis software supported by G.d.M.. S.S. performed experiments and contributed to analyzing the data. K.O. contributed to organoid culturing and performing experiments. S.W. contributed to the *Hydra* experiments. F.M., S.S., G.d.M., P.S. and P.L. wrote the manuscript.

## Data availability

Source data supporting the findings of this study are available from the corresponding authors on request.

## Code availability

Codes used to generate the findings of this study are available from the corresponding authors on request.

**Supplementary Video 1**

Maximum intensity projections of time lapse imaging of nine intestinal organoids expressing the Fucci2-reporter (hGem-mVenus and hCdt1-mCherry) corresponding to Fig. 1g. At the end of the movie, 3D-animations for selected organoids are shown.

**Supplementary Video 2**

Scan through individual planes in the z-direction comparing the image quality of detection objective 1 (left) and 2 (right) and the fused data (middle) corresponding to the intestinal organoid shown in Fig. 1 h.

**Supplementary Video 3**

Maximum intensity projections of time lapse imaging of the regeneration of three *Hydras* expressing ecto [ß-act::RFP] corresponding to Fig. 1i.

**Supplementary Video 4**

Scan through individual planes in the z-direction comparing the image quality of detection objective 1 (left) and 2 (right) and the fused data (middle) corresponding to *Hydra* shown in Fig. 1 j.

**Supplementary Video 5**

Maximum intensity projections of time lapse imaging of two liver organoids expressing mg-GFP and H2B-mCherry corresponding to Fig. 1k.

**Supplementary Video 6**

Maximum intensity projections of time lapse imaging the human colon cancer organoids expressing H2B-mNeon and STAR-sTom-NLS corresponding to Fig. 1l.

**Supplementary Video 7**

Maximum intensity projections of time lapse imaging of two parotid salivary gland organoids expressing H2B-mCherry corresponding to Extended Data 2l.

**Supplementary Video 8**

Maximum intensity projections of time lapse imaging of four gastruloids expressing H2B-mIRFP corresponding to Extended Data 2j.

**Supplementary Video 9**

Maximum intensity projections of time lapse imaging of intestinal organoids expressing the Fucci2-reporter (hGem-mVenus and hCdt1-mCherry) that were imaged in four different conditions (from left to right): control, NaCl, PGE and Forskolin.

**Supplementary Video 10**

Maximum intensity projections of time lapse imaging of intestinal organoids expressing H2B-mCherry.

**Supplementary Video 11**

Maximum intensity projections of long term time lapse imaging the human colon cancer organoids over almost 12 days expressing H2B-mNeon.

**Supplementary Video 12**

Scan through individual planes in the z-direction of gastruloids expressing Lck-GFP 42h, 66h and 90h after cell seeding corresponding to Fig 2h.

## Methods

### Microscope

The presented light sheet microscope consists of two illumination, two detection and a transmitted light beam paths. For illumination 60 mW 488 nm (488-60), 80 mW 515 nm (515-80), 50 mW 561 nm (561-50) and a 100 mW 638 nm (638-100, all lasers from Omicron-Laserage Laserprodukte) laser were used. Lasers were combined in a laser combiner (LightHUB+, Omicron-Laserage Laserprodukte) and coupled into a single-mode optical fiber with a 0.7mm collimated beam output..

The collimated illumination light from the fiber is first reflected by a mirror (BB05E02, Thorlabs) mounted in a kinematic mount (POLARIS-K05S2, Thorlabs) and passed through a filter wheel (FW212C, Thorlabs) containing neutral density filters (NE510B, NE520B, NE530B, Thorlabs) to further attenuate the intensity by a factor of 10, 100 or 1000.

The laser beam is reflected by a system of four galvanometric mirrors (6210H, Novanta Cambridge Technology) that are placed in custom made aluminum mounts. By a compound movement of the four scanners the beams can be translated and rotated on the image plane (XY) to generate light sheet by scanning as well as translated and rotated in YZ plane for focus adjustment. After passing through a scan lens made from two achromatic lenses (47-718, Edmund Optics) the illumination beam passes through a tube lens made from two achromatic lenses (49-281 and 49-283 Edmund Optics) followed by a custom triplet lens (plano-concave, r=41.67, 7980-0F, biconvex, r=49.87, S-TIH4, plano-concave, r=41.67, S-TIH53, Optimax) and a 10X water immersion objective with an NA of 0.2 (T Plan EPI SLWD 10X, Nikon) and a glass window. The illumination beam reaches the sample at an angle of 30° with the horizontal axis crossing an air glass and glass water interfaces. The custom designed correction triplet lens compensates for chromatic and spherical aberrations caused by different media in the light path to achieve diffraction limited resolution.

To switch between the two illumination beam paths, a D-shaped pickoff mirror (PFD10-03-P01, Thorlabs) placed at an angle of 45° degrees after the scan lens was used to reflect the beam either to the light path of illumination one or two. Both illumination paths contain a custom-made retroreflector system that contains two prism mirrors (15-599, Edmund Optics) mounted on a linear stage (UMR25, Newport) to adjust beam collimation at the back focal plane of the illumination objective and axial position of the illumination beam focus.

On the detection side, the fluorescence signal is collected by each of the two 16X water immersion objectives with an NA of 0.8 (CFI75 LWD 16X W, Nikon) and a working distance of 3 mm, which are opposing each other. The signal is reflected by two mirrors to match the 200mm focal length of the tube lens (T12-LT-1X, Nikon) and passes through motorized filter wheels (LEP filter wheel, Ludl) each equipped with the following emission filters FF01-515/LP-25, FF01-523/610-25, FF01-542/27-25 (all from Semrock) and ZET405/488/561/640mv2 (Chroma Technologies). The fluorescence images are acquired using ORCA-Fusion sCMOS cameras (C14440-20UP, Hamamatsu). Each camera is mounted together with the filter wheel on a custom aluminum mount that is placed on a manually adjustable liner stage (M-UMR5.25, Newport) to bring the two views in focus.

In one of the detection beam paths a LED light (LED770L, Thorlabs) is reflected by a 750 nm short-pass dichroic mirror (FF750-SDi02, Semrock) to the objective which serves as a condenser to acquire transmitted light images, while the emitted fluorescence light still passes through the dichroic mirror to the camera.

### Chamber fabrication and mounting

To mount the chambers into the microscope, individual thermoformed chambers are fixed in a 3D printed sample holder which has room for four chambers. This holder is placed onto a XYZ motorized sample positioning stage assembled by combining three piezo stages (2 x CLS 3232-S, 1x CLS 3282-S, SmarAct). This system allows a maximal travel distance across the chambers of ∼50 mm (long axis). Since water immersion objectives are used for detection, the chambers are lowered into a water reservoir with the objectives immersed below the water surface. The water reservoir and the sample stage are covered by a lid to minimize water evaporation. The sample handling area of the microscope is inside an incubator box to ensure an environmentally controlled area for temperature (CUBE2, Life Imaging Services) and CO2 (LS2 Live gas controller, Viventis Microscopy).

For the chamber fabrication process open top chambers in a multi-well format are produced from FEP (fluoroethylene propylene) using a vacuum thermoforming process. FEP-foil (Adtech Polymer Engineering) with a thickness of 127 μm was cut in quadratic sheets of approximately 15 x 15 cm. A sheet of foil was then clamped inside a vacuum forming machine, where the foil was heated up for 8 minutes. After the foil has heated up, custom made aluminum pieces are placed below the foil and serve as molds for the chambers. Under vacuum, the foil is formed around the molds for 30 seconds. As a last step, the chambers are cut out manually from the thermoformed FEP foil.

### Microscope control software and electronics

All parts of the microscope are controlled by a microscope control software (Viventis Microscopy). The controller and sensor module of the positioning system (MCS2-MOD, MCS2-S, SmarAct) and the driver of the galvanometric scanners (673, Novanta Cambridge Technology) were powered and connected according to manufacturer instructions in a custom electronics enclosure. Digital and analog signals to control lasers and galvanometric scanners were generated by an FPGA based real time controller (Viventis Microscopy).

### Microscope used for benchmarking experiments

For the comparison with the microscope presented in this work, a microscope system as described in [13], [17] with the following configuration was used: For excitation the light sheet microscope is equipped with a 488 nm (LuxXPlus 488-60), a 561 nm (OBIS 561-50) and a 630 nm (LuxXPlus 630-150) laser. For illumination two 10X water immersion objectives with an NA of 0.3 (CFI Plan Fluor 10XW, Nikon) are installed. The light sheet is generated by scanning the laser beam with a galvanometric scanner system and has a thicknesses (FWHM) of 2.2 μm approximately. A 25X 1.1 NA objective (CFI75 Apo 25XW; Nikon) is used to collect the emitted fluorescence. The images are acquired by an sCMOS camera (Zyla 4.1, Andor). Before the camera a filter wheel is placed offering the following filters: 488 LP Edge Basic Longpass Filter - F76-490; 561 LP Edge Basic Longpass Filter-F76-561; HC Dualband Emitter R 488/568 - F72-EY2, Semrock, AHF.

### Image Processing

Fusion of the data obtained from the two detection objectives is performed as follows. First, the stacks coming from the two detection objectives are registered to perfectly overlap by using the open-source Python imaging library DIPY. A rigid-body transformation is used to compensate for small mechanical misalignments between the two detection objectives.

Second, to fuse the data coming from the two detection objectives, the optimal z-plane is identified to switch from one view to the other to ensure the highest possible image quality in the fused image stack. Therefore, for both image stacks an image quality score is calculated plane by plane based on the Shannon entropy of the normalized discrete cosine transform (DCT) as discussed in [21]. The metric allows to set the switching z-plane to that point, where the opposing detection objective shows the higher image quality.

The stacks were then fused using a sigmoidal function centered at the switching plane and a constant intensity offset was subtracted to compensate for the background of the cameras

### Single cell tracking

Single cells of the intestinal organoid and the gastruloids were tracked manually by making use of the Fiji-Plugin Mastodon [30]. For the intestinal organoids, we used the numerical feature extraction of Mastodon to compute the mean intensity of all the spots marking the position of the cells. The mean intensities together with the spot positions for each track was exported as csv data table.

For the tracked cells of the gastruloid, we exported only the spot positions for each track as csv data table. The analysis of the cell tracks was done by custom written Python scripts relying on functions from open-source Python libraries numpy, pandas, seaborn and matplotlib.

The mean square displacement (MSD3D) for the trajectories ***r***_*i*_(*t*) of the gastruloid cells labelled with index *i* was calculated as follows [39]:

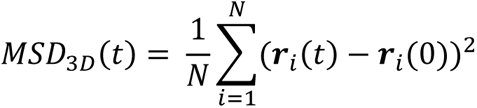

With *N* the number of cells and ***r***_*i*_(0) being the initial position of the cell.

### Single cell segmentation

Single cell segmentation of the gastruloids was performed with Cellpose 2.0 [29]. To train the model, a stack from a gastruloid 42h, 66h and 90h after cell seeding was used. Each plane of the stacks was manually annotated, and all three stacks were used to train a model for cell segmentation. To generate the masks in 3D we used the 2D prediction for image plane, which were then stitched together based on the overlap of the masks.

To extract features from the 3D-segmentation we relied on the 3D feature extraction from Python library scikit-image using the version 0.20.0.dev0. The extracted features were analysed with custom-written Python scripts based on the open-source libraries numpy, pandas, seaborn and matplotlib.

### Human material, mouse and stem cell lines

#### Human organoids

Patient-derived organoids identified by the HUB code P-19bT CRC organoids are cataloged at www.huborganoids.nl and can be requested at. Distribution to third (academic or commercial) parties has to be authorized by the Biobank Research Ethics Committee of the University Medical Center Utrecht (TCBio) at request of Hubrecht Organoid Technology (HUB). The movieSTAR, is based on a transposase-based integration method (movieSTAR: Tol2 insulator8xSTAR-min.pLGR5-sTomato-NLS-pA-PGK-H2BmNeonGreen-2A-Puro) as previously described [40].

#### Mouse Strains

All animal experiments were approved by the Basel Cantonal Veterinary Authorities and conducted in accordance with the Guide for Care and Use of Laboratory Animals. Male and female outbred mice between 7 and 15 weeks old were used for all experiments. The study employed the following mouse lines: B6/N x R26 Fucci2 (Tg/+) intestines, generously provided by J. Skotheim at Stanford University, to generate organoids. These organoids were subsequently infected with pGK Dest H2B-miRFP670 (Catalog no. 90237, Addgene). Additionally, (C57BL/6J) WT mice, R26-mG mice (#007676 [41]), obtained as a kind gift from T. Hiiragi at the Hubrecht Institute, R26-H2B-mCherry mice [42], received as a kind gift from C. Tsiairis at FMI Basel, and R26-mG/H2B-mCherry (heterozygous) mice, generated in-house following the procedure described in reference [43], were utilized. To monitor cell and nuclear shape, homozygous R26-mG mice were crossbred with homozygous R26-H2B-mCherry mice.

For genotyping of the newly generated R26-mG/H2B-mCherry mouse strain, the One Taq HS QuickLoad PCR reagent was used (Catalog no. M0488L, NEB). The genotyping process utilized the primer sets and cycling conditions listed in Supplementary Table 1.

#### Mouse stem cell lines

##### Cloning

As a nuclear marker H2B-miRFP was used. The protein coding sequence of H2B was amplified from the pCAG-H2BtdiRFP-IP plasmid obtained from Addgene (Catalog no.47884, Addgene) using KAPA HiFi polymerase (Catalog no. KK2501, Kapa Biosystems) introducing restriction sites for XhoI (Catalog no. R0146s, NEB) and SmaI (Catalog no. R0141S, NEB). These sites were subsequently used to introduce the PCR product into linearized pmiRFP670-N1 plasmid (Catalog no.79987, Addgene) using T4 DNA Ligase (Catalog no. M0202S, NEB). The protein coding sequence of H2B-miRFP670 was then amplified via PCR introducing attB sites to eventually clone the PCR product into pDONR221 (Catalog no. 12536017, Thermo Fisher) using Gateway BP Clonase II (Catalog no. 11789020, Thermo Fisher) creating an entry clone. Using LR Clonase II (Catalog no. 11791100, Thermo Fisher) the sequence of interest was introduced into an expression clone making use of pPB-UbC-DEST-pA-pgk-hph (kind gift from J. Betschinger, Novartis) creating the final PiggyBac construct containing H2B-miRFP670 driven by an Ubc promoter and a hygromycin selection cassette (hph) under a pgk promoter flanked by 5′ and 3′ repeats to allow PBase insertion.

As a membrane marker a Lck-tagged GFP was used. First, a pPB-UbC-DEST-pA-pgk-hph vector (kind gift from J. Betschinger, Novartis) based on plasmids published in [44] was subcloned to introduce a single MluI cloning site. In order to do so, the initial backbone was linearized with XbaI (Catalog no. R0145T, NEB) and BbsI (Catalog no. R0539S, NEB) and a gBlock (IDT) was used to introduce the MluI cut site via Gibson assembly (Catalog no. E2611L, NEB). The protein coding sequence of Lck-GFP was amplified from the Lck-GFP plasmid (Catalog no.61099, Addgene) using Phusion high fidelity polymerase (Catalog no. F530S, Thermo Fisher). Using MluI (Catalog no. R3198S, NEB) the expression vector was linearized, and Gibson assembly was used to incorporate the LcK-GFP sequence into the final PiggyBac vector.

For the above-described cloning efforts the Qiaquick gel extraction kit (Catalog no. 28706, Qiagen) was used to purify PCR and digestion products from agarose gels. Mix & Go E. Coli transformation kit (Catalog no. T3001, Zymo Research) treated DH5alpha cells (Catalog no. 18265-017, Invitrogen) were used for transformation and plasmid amplification. Bacterial selection was performed with Kanamycin (50 µg/ml) for Addgene plasmids #61099 and #79987 and Ampicillin (100 µ/ml) for all other plasmids. Plasmid purification was performed using QIAprep Spin Miniprep Kit (Catalog no. 27106, Qiagen) and NucleoBond Xtra Midi kit (Catalog no. 740410.50, Macherey-Nagel).

Engineering of monoclonal mouse embryonic stem cell lines:

Embryonic stem cell lines E14 and CGR8 (see below) were transfected using Lipofectamine2000 (Catalog no. 11668030, Thermo Fisher Scientific) to deliver the above mentioned PiggyBac plasmids. A total of 800 ng of purified DNA (400ng of expression vector and 400 ng of the vector carrying the PiggyBac transposase (kind gift form J. Betschinger, Novartis)) were diluted in 50 µl of OptiMEM (Catalog no. 31985062, Gibco)). In a separate tube 2 µl of Lipofectamine2000 was diluted in 50 µl. After 5 min of incubation at room temperature both mixtures were combined and thoroughly mixed before subsequent incubation at room temperature for 20 min. In the meantime, 2.5×10^5^ cells were seeded into a gelatine coated well of a 24well plate containing Serum medium. Serum medium consists of GMEM (Catalog no.G5154, Sigma) supplemented with 10% v/v ESC-grade FBS (Catalog no. 16141079, Invitogen), 1x GlutaMAX (Catalog no. 35050038, Invitrogen), 1x MEM-NEAA (Catalog no. M7145-100ML, Sigma), 1x sodium pyruvate (Catalog no. 11360070, Invitrogen), 1x β-Mercaptoethanol (Catalog no. 21-985-023, Fisher Scientific), 3 μM CHIR99021 (Chir) (Catalog no. 72054, Stem Cell Technologies), 1 μM PD0305901 (Catalog no. 100-0248, Stem Cell Technologies) and 0.01 μg/ml LIF (Catalog no. 78056, Stem Cell Technologies). While the cells were still in suspension the transfection mix was added and the plate was agitated in order to equally distribute the solution. Now the plate was incubated over night before the medium was replaced with fresh Serum medium. After 3 days, antibiotic selection with hygromycin B (Catalog no. 10-687-010, Fisher Scientific) (200 µg/ml) was started and continued for >7 days. Finally single cells were sorted into a gelatin coated 96well plate containing Serum medium supplemented with Pen/Strep (Catalog no. 15140122, Gibco). Based on fluorescence and morphology, monoclonal colonies were selected and expanded.

Embryonic stem cell lines CGR8-Lck-GFP-H2B-mCherry and E14-H2B-mCherry were generated from their parental lines CGR8 and E14 respectively. Both E14 and CGR8 cell lines are of 129P background and were kind gifts from M. Lutolf (IHB Roche). Cells were tested routinely for mycoplasma via PCR.

#### Sample preparation

##### Parotid salivary gland organoids

Parotid salivary glands were dissected following previously established protocols [45]. After dissection, the gland tissue was cut into small fragments using a razor blade. Thereafter, fragments were enzymatically digested with Trypsin 0.05% EDTA for 10-15 min. Cells were passed through a 40um strainer, and embedded in 50% Matrigel (for organoid culture, Catalog no. 356255, Corning) and supplied with Basal culture medium consisting of DMEM/F12 (with 15 mM HEPES) (Catalog no. 36254, Stem Cell Technologies), 1x GlutaMAX (Catalog no. 35050038, Thermo Fisher Scientific)), and 100 μg/mL Pen/Strep (Catalog no. 15140-122, Gibco)) with 0.5 nM Wnt (NGS) (Catalog no. N001, UproteinExpress), 1 µg/ml Recombinant R-Spondin1 (kind gift from Novartis), 100 ng/ml Recombinant Noggin (Catalog no. 250-38, PrepoTech), 1x B27 supplement (Catalog no. 17504044, Thermo Fisher Scientific), 1.25 mM NAC (Catalog no. A7250-100G, Sigma Aldrich), 50 ng/ml hEGF (Catalog no. 236-EG-200, RnD Systems), 10 ng/ml hNRG1 (Catalog no.5898-NR-050, RnD Systems), 0.5 µM A83-01 (Catalog no. 2939, Tocris), 100 ng/ml hIGF1 (Catalog no.291-G1-200, RnD Systems), 50 ng/ml hFGF2 (Catalog no. 233-FB-025, RnD Systems), and 10 µM ROCK-Inhibitor (Catalog no. Y-27632, Stem Cell Technologies). Organoids were maintained by hard splitting every week. Directly after the initial hard-split the culture medium composition changed to 500 ng/ml Recombinant R-Spondin1, 100 ng/ml Recombinant Noggin, 1x B27 supplement, 1.25 mM NAC, 1 ng/ml hNRG1, 100 ng/ml hIGF1, 50 ng/ml hFGF2, and 100 μg/ml Primocin (Catalog no. ant-pm-1, Invivogen). Samples on the light-sheet were imaged 7 days after a hard-split.

##### Human colon cancer organoids

Human CRC organoids were maintained in 70% Matrigel-drops (for organoid culture, Catalog no. 356255, Corning). The medium composition for the patient-derived tumor organoid was: Basal culture medium (see parotid salivary gland organoid section) supplemented with 0.5 µg/mL recombinant R-Spondin1 (kind gift from Novartis), 100 ng/mL recombinant Noggin (Catalog no. 250-38, PrepoTech), 1x B27 supplement (Catalog no. 17504044, Thermo Fischer Scientific), 1.25 mM NAC (Catalog no. A7250-100G, Sigma Aldrich), 50 ng/mL hEGF (Catalog no. 236-EG-200, RnD Systems), 100 ng/mL hIGF1 (Catalog no. 291-G1-200, RnD Systems), 50 ng/mL hFGF2 (Catalog no. 233-FB-025, RnD Systems), 10 nM Gastrin (Catalog no. G9145-.1MG, Sigma Aldrich), 500 nM A83-01 (Catalog no. 2939, Tocris), 3 uM SB202190 (Catalog no. 1264, Tocris), and 100 μg/ml Primocin (Catalog no. ant-pm-1, Invivogen). Preceding imaging using the light-sheet microscope setup, organoids were dissociated into individual cells and then embedded into 60% Matrigel drops within a specialized holder designed for light-sheet imaging experiments. The culture medium, mentioned earlier, was supplemented with 10 µM ROCK-Inhibitor (Catalog no. Y-27632, Stem Cell Technologies) during the first 4-days. Subsequently, medium renewal occurred every 48 hours without the ROCK-Inhibitor.

##### Hepatic organoids

The establishment of murine hepatic organoid culture was carried out with some adaptations as described in [46]. In brief, a hepatectomy was performed on a mouse from the above described R26-mG/H2B-mCherry strain (see mouse strain section above). After thoroughly washing the explanted liver in ice cold PBS the tissue was minced using surgical scissors. After a fine tissue paste formed, the material was resuspended in 50 ml of ice cold DMEM (high glucose, Catalog no. D5796-500ML, Sigma Aldrich) and transferred into a 50 ml tissue culture tube. After inverting the tube multiple times, the material was sedimented via centrifugation (80 g at 4°C for 2 min). This step was repeated 3 times. Now the supernatant was decanted, with around 10 ml remaining. The tissue fragments were then further dissociated by extensive pipetting with a 5 ml serological pipette (pipetted up and down at least 30 times). Centrifugation (200 g at 4°C for 5 min) was used to pellet the dissociated tissue before the pellet was resuspended in 15 ml prewarmed digestion mixture (DMEM high glucose supplemented with collagenase (Catalog no. C9407-25MG, Sigma Aldrich) at a final concentration of 0.25 mg/ml. After one more round of centrifugation the pellet was resuspended in 45 ml of prewarmed digestion mixture and subsequently incubated at 37°C with moderate shaking for 45 min. After 45 min an aliquot of the suspension was assessed via microscopy to confirm the presence of ductal structures. After ductal structures were found the digestion was terminated by pelleting the supernatant (centrifugation at 4°C and 200 g for 5 min). The pellet was subsequently resuspended in wash medium and centrifuged again with the same settings. Finally, the material was resuspended in 50 % Matrigel (for organoid culture, Catalog no. 356255, Corning) and Hepaticult (Catalog no. 06030, Stem cell technologies) and plated in 50 µl droplets in a 12well tissue culture plate. After the Matrigel drops were solidified 1 ml of Hepaticult medium was added to each well. Medium was changed when passaging the organoids. Organoids were passaged at least 5 times before starting light sheet acquisitions with the newly established organoid line. For light sheet sample mounting 10 µl droplets of mechanically split organoids resuspended in 50 % Matrigel were seeded into a sample holder with a diameter of 1 mm. After solidifying for 15 min at 37°C the Matrigel droplets were covered with 300 µl of Hepaticult.

##### Intestinal organoids

Mouse small intestinal organoids were established and cultured as previously described in [13]. Intestinal organoids expressing the Fucci2-reporter (see mouse strain section above) were previously established as described in [13], [17], cultured in droplets of 50% Matrigel (for organoid culture, Catalog no. 356255, Corning) with IntestiCult Organoid Growth Medium (Catalog no. 06005, Stem cell technologies) and were kept in IntestiCult OGM supplemented with 100 μg/ml Pen/Strep (Catalog no. 15140122, Gibco) for maintenance. For light sheet experiments, organoids were collected 5 days after mechanical disruption. Organoids were embedded in Matrigel and ENR medium (1:1 ratio) in custom FEP-chambers (see Extended data Fig. 1b). ENR medium is composed of advanced DMEM/F-12 with 15mM HEPES (Catalog no. 36254, Stem Cell Technologies) supplemented with 100 μg/ml Pen/Strep (Catalog no. 15140-122, Gibco), 1× Glutamax (Catalog no. 35050061, Gibco), 1× B27 (Catalog no. 17504044, Gibco), 1x N2 (Catalog no. 17502048, Thermo Fisher Scientific), 1 mM NAC (Catalog no. A7250-100G, Sigma Aldrich), 500 ng/ml R-Spondin (kind gift from Novartis), 100 ng/ml Noggin (Catalog no. 250-38, PrepoTech) and 100 ng/ml murine EGF (Catalog no. 315-09-100ug, RnD Systems). After 20 min of solidification at 37°C, 300 µl of ENR medium supplemented with 20 % Wnt3a-conditioned medium (Wnt3a-CM) was added. Perturbation studies on mouse small intestinal organoids were performed in ENR medium supplemented with 0.5 μM PGE, 5 μM Forskolin, 250 mM NaCl solution, or DMSO as vehicle control.

##### Hydra

Light sheet imaging of regenerating *Hydra* was performed on the *ecto[β-act::RFP]/endo[β-act::GFP]* “*Reverse Watermelon”* line of *Hydra vulgaris* [47]. *Hydra* culture was maintained at 18°C in Volvic mineral water. Animals were fed three times per week with freshly hatched *Artemia nauplii*.

For the light sheet acquisition, adult, non-budding animals were used. Animals were transferred to *Hydra* medium (1 mM CaCl2, 0.2 mM NaHCO3, 0.02 mM KCl, 0.02 mM MgCl2, and 0.2 mM Tris-HCl (pH 7.4)) and spheroids were cut as previously described [48]. In short, the animals were bisected with the initial cut being directly under the tentacle ring. Two tissue rings were obtained from each animal by sequentially cutting the body axis. The rings were split in two to three rectangular pieces. The tissue fragments were left to fold for 4 hours in dissociation medium (3.6 mM KCl, 6 mM CaCl2, 1.2 mM MgSO4, 6 mM sodium citrate, 6 mM sodium pyruvate, 4 mM glucose, and 12.5 mM N-tris(hydroxymethyl)methyl-2-aminoethanesulfonic acid (pH 6.9)) at room temperature. Properly closed spheroids of a typical size (300-500 μm in diameter) were selected for the imaging. The *Hydra* were imaged in sample chambers that were filled with 500 µl *Hydra* medium.

##### Gastruloids

Gastruloids used in this study have been prepared as previously described [26]. In brief mouse embryonic stem cells (ESC) were maintained on gelatin-coated culture plates (6-well) in N2B27 medium consisting of 50 % DMEM/F12 (Catalog no. 21331020, Gibco) and 50 % Neurobasal medium (Catalog no. 21103049, Gibco) supplemented with 1x N2 (homemade) and 1x B27 serum free supplement (Catalog no. 17504044, Gibco), 1x GlutaMAX (Catalog no. 35050061, Gibco), HEPES (Catalog no. 83264, Sigma Aldrich) and 1x β-mercaptoethanol (Catalog no. 21-985-023, Gibco). For stem cell maintenance N2B27 medium was supplemented with 3 μM CHIR99021 (Chir) (Catalog no. 72054, Stem Cell Technologies), 1 μM PD0305901 (Catalog no. 100-0248, Stem Cell Technologies) and 0.01 μg/ml LIF (Catalog no. 78056, Stem Cell Technologies). ESCs were split every other day by disassociating colonies using 400 µl of Accutase (Catalog no. A6964-500ML, Sigma Aldrich). After visual inspection 2 ml of wash medium (DMEM/F12 with 0.1% BSA Catalog no. 15260037, Thermo Fisher Scientific) was added to create a cell suspension. Cells were pelleted via centrifugation (300 g, 4°C, 5 min) and resuspended in N2B27 medium. This step was repeated one more time to remove remaining traces of compounds added to maintain naïve pluripotency. Cells were usually split in a ratio between 1:10 and 1:15. An aliquot of the same cell suspension was used for cell counting (TC20, Biorad).

40 µl of N2B27 containing 300 ESCs were seeded into each well of a 96well ultra-low attachment U-bottom plate (Catalog no. 7007, Corning). After 48 hours, the formed aggregates were pulsed with 150 µl of N2B27 supplemented with 3 μM Chir for 24 hours. After a total 72 hours post seeding, 140 µl of N2B27 was removed and 160 µl of fresh N2B27 medium was added. At 96 hours post seeding a last medium change was performed removing 150 µl of medium and replacing it with an equal amount of fresh N2B27 medium. Gastruloid culture was not performed for longer than 120 hours. For light sheet imaging gastruloids have been collected at either 42, 66, 90 or 96 hours post seeding. Gastruloids have been mounted into pockets of custom thermoformed sample holders (Extended Data Fig. 2c) holding 500 µl of N2B27 for suspension culture. For indicated cases the pockets were covert with 40 % of Matrigel (for organoid culture, Catalog no. 356255, Corning) diluted in N2B27 to ensure minimal mechanical rotations of forming gastruloids. For the 42 and 66 h timepoints 5 % of fluorescent cells were mixed with 95 % of their respective non fluorescent parental line. For later timepoints 10 % of fluorescent cells were used.

#### Endpoint immunofluorescence/DRAQ5staining

##### Small intestinal organoids

After live imaging of murine intestinal organoids expressing the Fucci2-reporter (see above), the sample holder was removed from the microscope in order to perform sample fixation using 4 % PFA (Catalog no. 15714, Electron Microscopy Sciences) and 0.08 % glutaraldehyde (Catalog no. 16019, Electron Microscopy Sciences) diluted in PBS for 25 min at room temperature. Next, samples were washed with PBS 3 times with 10 min incubations. In order to reduce auto fluorescent background potentially resulting from the use of glutaraldehyde a quenching step using sodium borohydrate (Catalog no. 452882-5G, Sigma Aldrich) (0.01 g/10 ml of PBS) was performed for 10 minutes at room temperature. Now the samples were permeabilized and blocked by incubating with 3 % donkey serum (Catalog no. D9663-10ML, Sigma Aldrich) and 2 % TritonX (Catalog no. T9284, Sigma Aldrich) in PBS for 1 hour. After permeabilization and blocking primary antibody solution using sheep anti Dll1 (Catalog no. AF3970, RnD Systems) and rabbit anti Lysozyme (Catalog no. A0099, Dako) antibodies at a concentration of 1:100 in PBS supplemented with 3 % donkey serum and 0.1 % TritonX was added. Primary antibody solution was incubated for 2x over night at 4°C before the sample was washed for 3×30 min with PBS at room temperature. Now, the samples were incubated with Fab fragments at a concentration of 1:250 in PBS supplemented with 3 % donkey serum and 0.1 % TritonX overnight. Donkey anti rabbit Fab fragments conjugated to Alexa 647 and donkey anti goat Fab fragments conjugated to Alexa 488 fluorophores were used (Catalog no. 705-607-003 and 711-547-003, Jackson Immuno Research). After washing the samples with PBS for 3×30 min at room temperature, the sample holder was remounted onto the light sheet microscope for imaging.

##### Gastruloids

Gastruloids were fixed in 4% PFA (Catalog no. 15714, Electron Microscopy Sciences) in PBS for 30 min at room temperature. After extensive washes with PBS, gastruloids were permeabilized with 1% Triton X (Catalog no. T9284, Sigma Aldrich) in PBS for 1 hour before an overnight incubation with DRAQ5 (Catalog no. 62251, Thermo Fisher Scientific) at a final dilution of 1:300 in PBS. After further extensive washing with PBS, gastuloids were mounted for light sheet imaging.

---

**Extended Data Fig 1.**
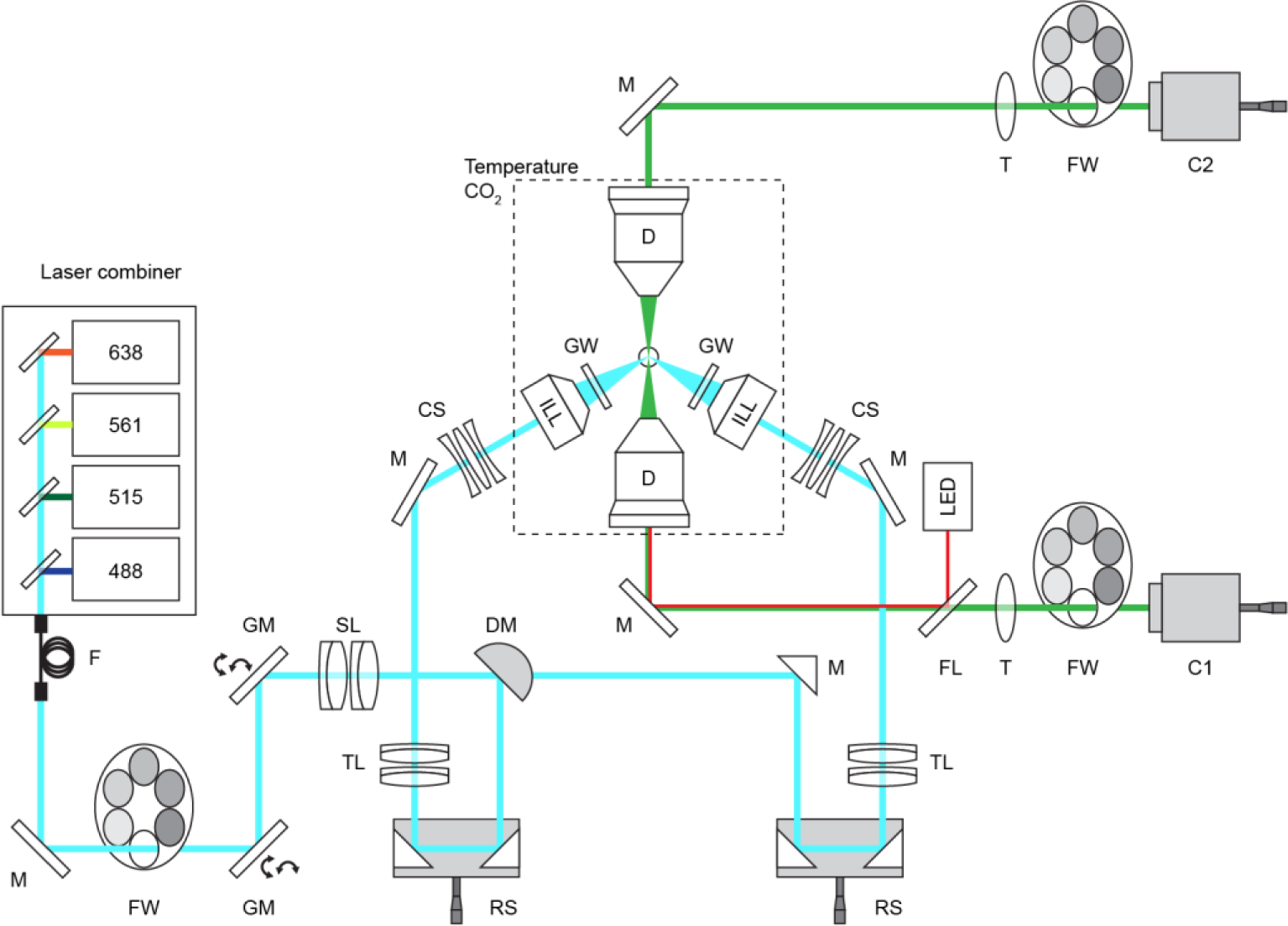
Light path of the dual view light sheet microscope. Schematic representation of the light path of the dual view and dual illumination light sheet microscope: fiber (F), mirror (M), filter wheel (FW), galvanometric mirror (GM), scan lens (SL), d-shaped mirror (DSM), reflector system (RS), tube lens (T), correction system (CS), illumination objective (ILL), glass window (GW), detection objective (D), dichroic mirror (DM), camera 1 and 2 (C1,C2). See Methods section.

**Extended Data Fig 2.**
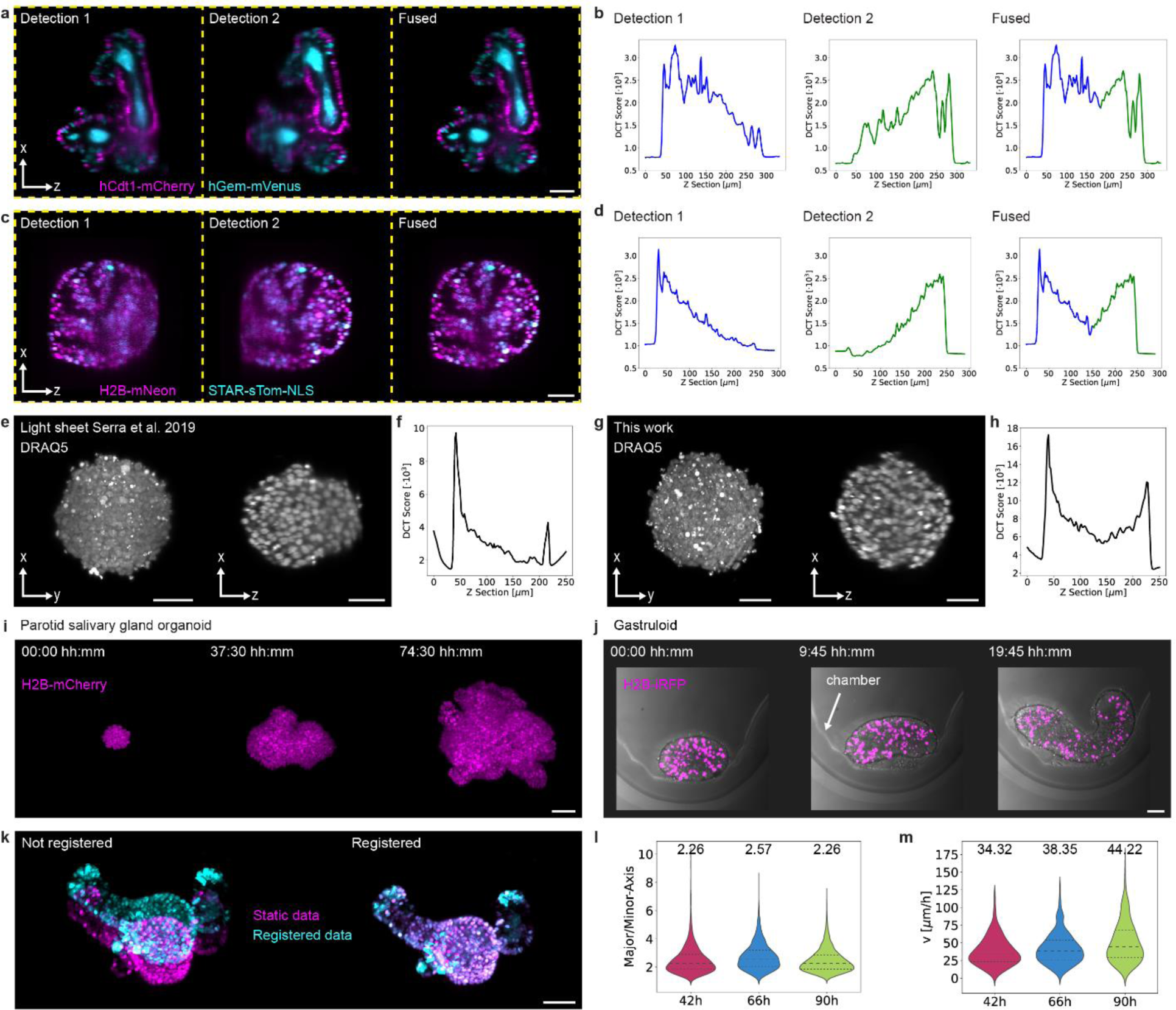
Comparison of the image quality using different light sheet microscopes, showcase acquisitions, an example of image registration and results of the single cell analysis of gastruloids in suspension. a) Cross section in XZ plane of the intestinal organoid shown in Fig 1f using Detection 1, Detection 2 and the fused data from both objectives. Scale bar 50 µm. b) Comparison of the image quality using the Shannon Entropy of the Discrete Cosine Transform (DCT) as metric for Detection 1, Detection 2 and the fused data. The DCT is calculated for each z section of the image stack corresponding to a. c) Cross section in XZ plane of the human colon cancer organoid shown in Fig 1l using Detection 1, Detection 2 and the fused data from both objectives. Scale bar 50 µm. d) Comparison of the image quality using the Shannon Entropy of the DCT as metric for Detection 1, Detection 2 and the fused data. The DCT is calculated for each z section of the image stack corresponding to c. e) Maximum intensity projection (MIP) along the Z axis and a cross section in the XZ plane of a gastruloid stained with the nuclear marker DRAQ5 imaged with the light sheet microscope published in [13]. Scale bar 50 µm. f) Image quality plot using Shannon Entropy of the DCT along individual z sections of the image stack corresponding to e. g) Maximum intensity projection (MIP) along the Z axis and a cross section in the XZ plane of a gastruloid stained with the nuclear marker DRAQ5 imaged with the light sheet microscope presented in this work. Scale bar 50 µm. h) Image quality plot using Shannon Entropy of the DCT along individual z sections of the image stack corresponding to g. i) MIPs of a parotid salivary gland organoid expressing H2B-mCherry at three different time points of a time lapse acquisition for around 3 days. Scale bar 50 µm. j) MIPs of a gastruloid expressing H2B-iRFP at three different time points of a time lapse acquisition over 20 hours shown together with transmitted light. Scale bar 50 µm. k) Example of the registration of an organoid, that was additionally fixed and stained after live imaging. Left image shows the overlay of the MIPs before registration and the right image shows the results after registration. Magenta shows the MIP of the last acquired timepoint and cyan shows the MIP after fixation and staining. Scale bar 50 µm. l) Comparison of the ratio of major and minor axis for the three different time windows investigated in gastruloids imaged in suspension. Measurements were performed over 33 timepoints on the 3D volumes of 3 gastruloids per imaging window (42 h, 66 h or 90 h). Summing up individual timepoints, a total of 3753 datapoints (42 h), 3340 datapoints (66 h) and 8861 datapoints (90 h) were analyzed. Median (values in figure), first and third quartile are shown. m) Violin plot of the velocity of cells imaged at 3 different windows of gastruloid development showing the median (values in figure) and the first and third quartile. The following numbers of datapoints (from 3 individual gastruloids) were tracked per observation window: n = 622 (42 h), n = 562 (66 h), n = 539 (90 h). Gastruloids were imaged in suspension.

**Extended Data Fig 3.**
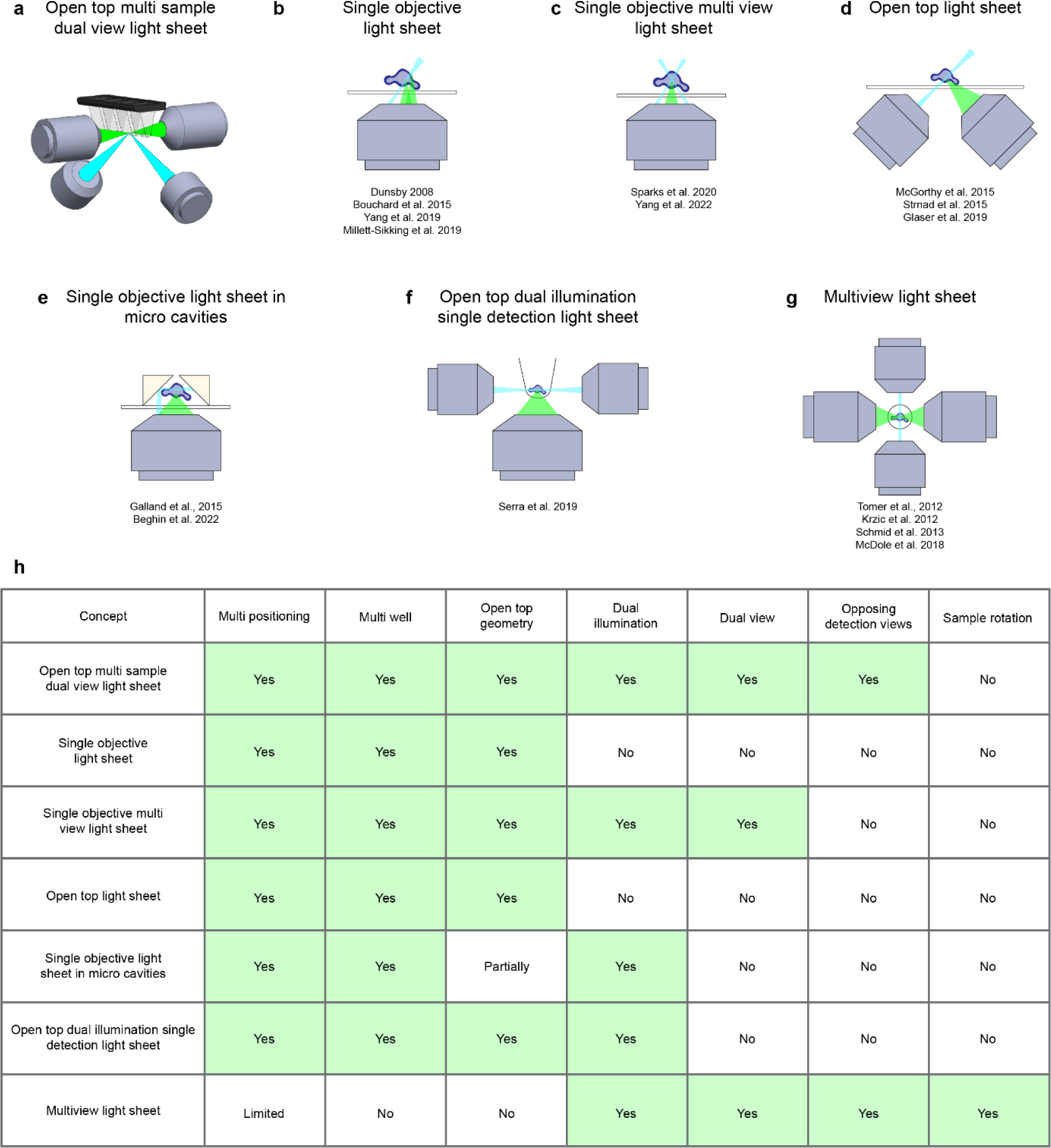
Schemes and comparison of key parameters among state-of-art light sheet microscopy methods. a) Scheme of the open top multi sample dual view light sheet microscopy (presented in this work). b) Scheme of single objective light sheet microscopy [10], [11], [33], [34]. c) Scheme of single objective multi view light sheet microscopy [35], [36]. d) Scheme of open top light sheet microscopy [9], [12], [37]. e) Scheme of single objective light sheet microscopy in micro cavities [32], [49]. f) Scheme of open top dual illumination single detection light sheet microscopy [13]. g) Scheme of Multiview light sheet microscopy [4]–[6], [38]. h) Comparison of key aspects of different state-of-art light sheet microscopy methods.

## Supplementary Information

**Supplementary Table 1:** Imaging settings used for all experiments.

| Figure | Sample | Fluorescent label | Excitation [nm] | Exposure Time [ms] | Z-Sections [2 μm spacing] | Time Interval [mins] | Acquisition time [h] |
| --- | --- | --- | --- | --- | --- | --- | --- |
| **1g, 2a-c, Ext Dat. 1a, Supp Video 1** | Intestinal organoids | hGem-mVenus<br>hCdt1-mCherry | 515<br>561 | 30<br>10 | 181 X<br>2μm | 10 | 67,3 |
| **1 l, Supp. Video 3** | *Hydra* | ecto[[β-act::RFP] | 561 | 10 | 801 X<br>2μm | 10 | 67,16 |
| **1 k, Supp Video 5** | Liver organoids | mg-GFP<br>H2B-mCherry | 488<br>561 | 10<br>10 | 201 X<br>2μm | 10 | 31,3 |
| **1 l, Supp. Video 6** | Human colon cancer organoids | H2B-mNeon<br>STAR-sTom-NLS | 488<br>561 | 10<br>10 | 201 X<br>2μm | 30 | 137 |
| **2 h,l** | Gastruloids | Lck-GFP | 488 | 10 | 350 X<br>1μm | 10 | 5,5 |
| **Ext. Dat. 2b, Supp Video 7** | Parotid salivary gland organoids | H2B-mCherry | 561 | 10 | 201 X<br>2μm | 30 | 74,5 |
| **Ext. Dat. 2c, Supp Video 8** | Gastruloids | H2B-iRFP | 638 | 10 | 251 X<br>2μm | 15 | 13,5 |
| **Supp. Video 9** | Intestinal organoids | mg-GFP<br>H2B-mCherry | 488<br>561 | 10<br>10 | 121 X<br>2μm | 3 | 4,9 |
| **Supp. Video 11** | Human colon cancer organoids | H2B-mNeon | 561 | 100 | 201 X<br>2μm | 30 | 280,5 |
| **Supp. Video 10** | Intestinal organoids | H2B-mCherry | 561 | 10 | 181 X<br>2μm | 10 | 21,5 |

**Supplementary Table 2:**
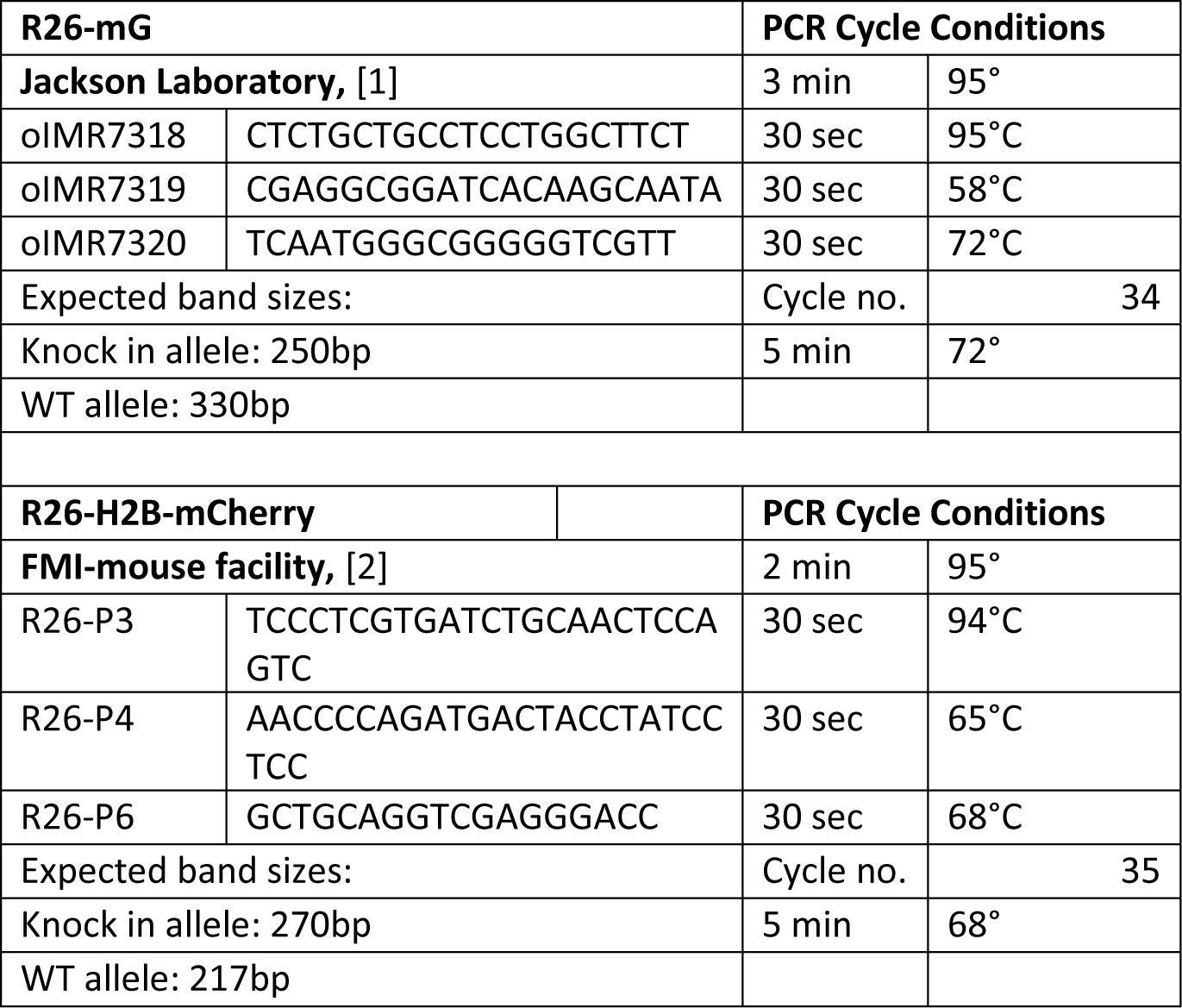
Primer sets and cycling conditions used for genotyping the newly generated R26-mG/H2B-mCherry mouse strain.

